# High-throughput production of functional prototissues capable of producing NO for vasodilation

**DOI:** 10.1101/2021.07.22.453323

**Authors:** Xiangxiang Zhang, Chao Li, Fukai Liu, Wei Mu, Yongshuo Ren, Boyu Yang, Xiaojun Han

## Abstract

Bottom-up synthesis of prototissues helps us to understand the internal cellular communications in the natural tissues and their functions, as well as to improve or repair the damaged tissues. The existed prototissues are rarely used to improve the function of living tissues. We demonstrated a methodology to produce spatially programmable prototissues based on the magneto-Archimedes effect in a high-throughput manner. More than 2000 prototissues are produced once within 2 hours. Two-component and three-component spatial coded prototissues are fabricated by varying the addition giant unilamellar vesicles (GUVs) order/number, and the magnetic field distributions. Two-step and three-step signal communications in the prototissues are realized using cascade enzyme reactions. More importantly, the two-component prototissues capable of producing nitric oxide (NO) cause vasodilation of rat blood vessels in the presence of glucose and hydroxyurea. The tension force decreases 2.59 g, meanwhile the blood vessel relaxation is of 31.2%. Our works pave the path to fabricate complicated programmable prototissues, and hold great potential in tissue transplantation in the biomedical field.

## Introduction

Natural tissues composed of multi-cellular components possess spatial hierarchical structures and collective behaviors triggered by inter communications among cells^1,2^. Bottom-up fabrication of prototissues is beneficial to understanding the interaction mechanism among cells in the tissues, as well as holding great potential in the field of biomedical engineering^3,4,5^.

The bottom-up assembled prototissues are roughly classified into two categories, i.e., randomly distributed and spatially well-defined structures. The randomly distributed prototissues were usually formed via noncovalent and covalent bonds. The prototissues composed of giant unilamellar vesicles (GUVs) were fabricated via electrostatic interactions^6^, while the prototissues composed proteinsomes were formed via alkyne-azide cycloaddition reaction^7,8^. The spatially well-defined prototissues were produced via external forces, including optical^9^, acoustic^10, 11, 12^, magnetic forces^13^. Optical tweezers were used to precisely build GUV colonies with various architectures of 2D (trigonal, square, pentagonal) and 3D (tetrahedral, square-pyramidal and three-layered pyramid) structures^9^. Acoustic fields were used to assemble various GUVs colony arrays, heterogeneous GUV/natural cells, as well as dynamic colony arrays^11^. In our previous work, diverse spatial programmed GUVs prototissues were achieved based on the magneto-Archimedes effect^13^. In addition, 3D bioprinting technique was able to precisely code more building blocks into higher-order prototissues^14,15,16^. The economic high-throughput method is still on demand to produce spatially coded prototissues.

Prototissues are not the simple aggregation of the protocells. The inter communications among protocells and the functions are more important for their biomedical applications. The signal communications in prototissues have been achieved^4,5^. The two-step cascade enzyme reaction was often used for this purpose. The hydrogen peroxide (produced in one component by glucose and glucose oxidase) transferred into other component containing horseradish peroxidase (HRP) to oxidize Amplex red into resorufin molecules in two-component prototissues^17,18^. The electric communication was realized in the 3D printing prototissues via light triggered hemolysin expression^19,20^. The three-step cascade reaction among the components or more complicated network in the prototissues was rarely reported^21^. The functions of prototissues were demonstrated with thermal-responding reversible contractions and expansions^8^, non-equilibrium biochemical sensing^22^, and collective deformation^14^. There is no report to improve the real tissue function using prototissues.

Herein, a versatile method was demonstrated to fabricate high-throughput spatial programmable prototissues based on the magnetic Archimedes effect. Thousands of prototissues with spatially controllable structures were made in one-go within 2 hours. The signal communications were investigated both in two- and three-component spatially coded prototissues through enzyme cascade reactions. More importantly, we demonstrated the prototissues were able to cause blood vessel expansion due to the NO produced from prototissues triggered by blood glucose. The versatile methodology proposed in this work provide a new path to fabricate complicated functional prototissues, which hold great potential in the biomedical field.

## Results and Discussion

### Generation of prototissues using Magneto-Archimedes effect

The spatial programmable prototissues were fabricated using the magneto-Archimedes effect in a home-made device (Fig. 1a and Supplementary Fig. 1) containing a nickel mesh (NM) inside petri dish on the top of a circular permanent magnet. The diamagnetic materials move to the weak magnetic field area in a nonhomogeneous magnetic field. This phenomenon is called magneto-Archimedes effect. Giant unilamellar vesicles (GUVs) are diamagnetic materials, which are used as the building blocks for prototissues. The magnetic potential energy **U(r)** of a GUV with radius **R** in the space magnetic field **H(r)** is given by formula (**1**)^23^.

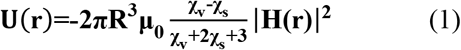

where **χ**_**v**_, **χ**_**s**_ and **μ**_**0**_ are the susceptibility of the GUV, the paramagnetic solution, and the permeability of the free space, respectively. Here, the magnetic potential energy **U(r)** is positive because **χ**_**v**_ is less than **χ**_**s**_. Magnetic force drives GUVs to the regions with lower magnetic field strength, according to the energy minimum principle. Therefore, the spatial structure of the prototissues can be predicted according to the distribution of magnetic field. The NM exhibits a strong magnetic response and causes a gradient magnetic field inside NM (Fig. 1b, bottom layer) and 140 μm above NM (Fig. 1b, top layer), where the blue areas indicate weak magnetic field regions. The weak magnetic field regions appear close to the wires in each grid in the NM surface (Fig. 1b, bottom layer), but in the center inside each grid 140 μm above NM surface (Fig. 1b, top layer). The NM was made with nickel wires (d = 60 μm). Each grid was 210 μm in square.

**Fig. 1.**
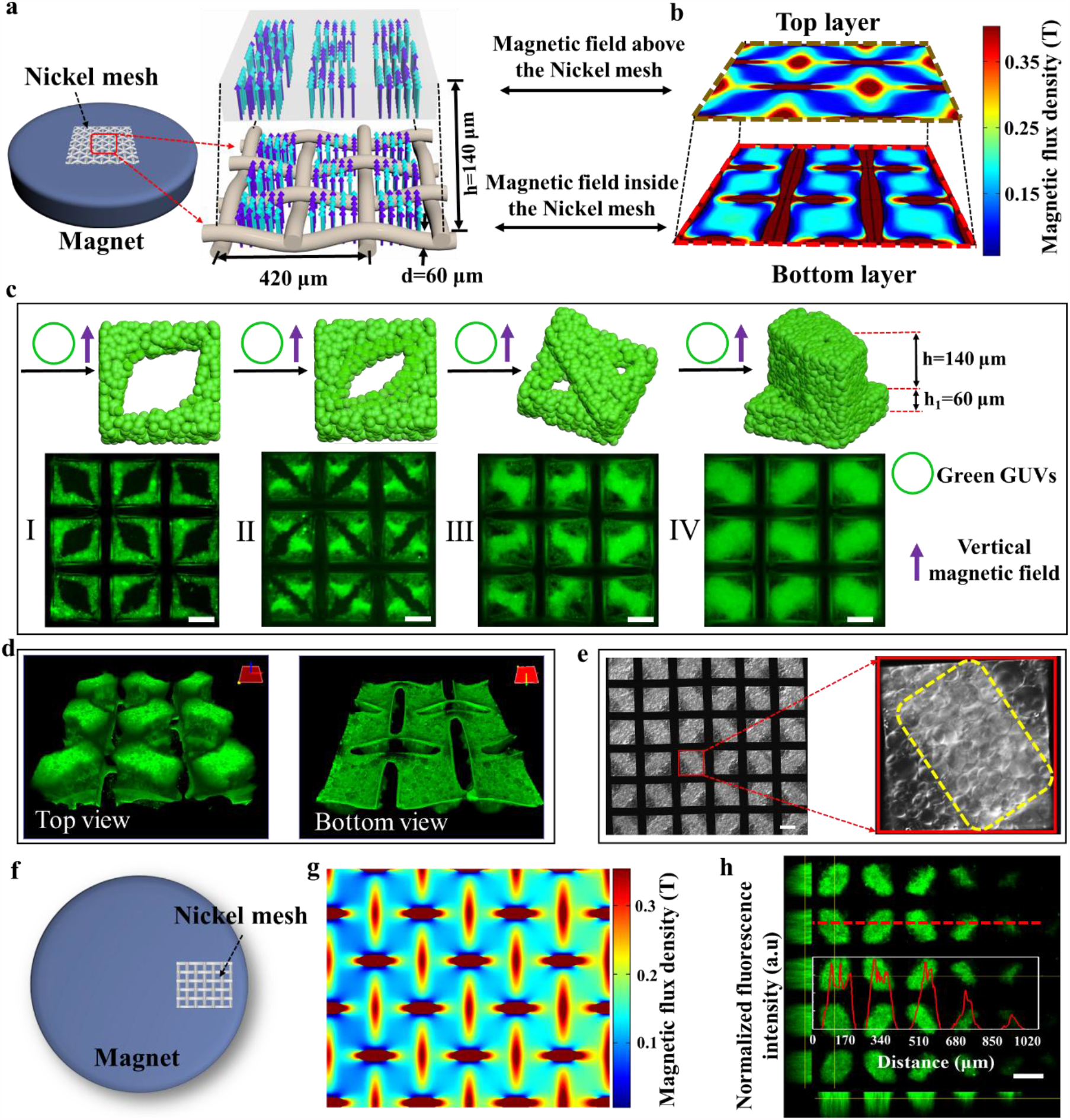
Generation of prototissues using Magneto-Archimedes effect. **a)** Schematic of the device with a woven nickel mesh (NM) at the center of a circular magnet. Cyan and blue arrows represented the strong and weak magnetic field, respectively. **b**) Simulation results of the magnetic field of the bottom layer and top layer, which indicated the distribution of the magnetic field inside and above the NM, respectively. Blue areas indicated the weak magnetic field regions. **c**) Schematic and fluorescence images of the prototissues assembled in a vertical magnetic field. With the number of added green giant unilamellar vesicles (gGUVs) increasing, the prototissues changed from single layer (I, II) to double layers (III, IV). **d)** 3D reconstructed confocal fluorescence images of the prototissue array with ‘protruded structure’ from top (left) and bottom (right) views. **e)** A differential interference contrast (DIC) image of the 3D prototissue array. The lighter region in the image indicated the protruding top layer, which was marked by the yellow dashed box in the right image. **f)** Schematic of the NM at the edge region of the circular magnet. **g)** The simulation result of the magnetic field of the NM in **(f). h)** A fluorescence image of gGUVs prototissues in the magnetic field **(g)**. Red line in the inset image corresponded to dashed line intensity analysis. The scale bars were 100 μm.

As expected, the GUVs were firstly assembled in the weak magnetic field regions (blue areas in Fig. 1b, bottom layer) inside each grid (Fig. 1c I). With the number of added GUVs increasing, GUVs gradually filled the space inside each grid of the NM to generate 60 μm thick bottom layer (Fig. 1c II) and further protruded to form 140 μm thick brick-shape top layer above the GUVs aggregations inside each grid (Fig. 1c III, IV), which were consisted with simulated field distribution (blue areas in Fig. 1b, top layer). The concentrations of the GUVs for fabricating the prototissues in Fig. 1c were 3×10^5^/mL for I, 5×10^5^/mL for II, 8×10^5^/mL for III and 1.2×10^6^/mL for IV, respectively. The 3D reconstructed confocal fluorescence images clearly showed the two-layer structures of GUV aggregates, containing brick-shape top layer (Fig. 1d, left image) and square bottom layer (Fig. 1d, right image). The differential interference contrast (DIC) microscope image further demonstrated the two-layer structure arrays in the NM (Fig. 1e), where the brighter regions indicated the protruded brick-shape layer marked by a yellow dashed box (Fig. 1e, right image). We named this structure as the “protruded structure”.

This prototissues fabrication method is easily scaled up to generate more than 2000 prototissues one time by using bigger NM (Supplementary Fig. 2). The diversity of this method is also demonstrated with producing various size of prototissues by using various NM with different length of the grid (Supplementary Fig. 3). In contrast, in the absence of a magnetic field, GUVs sank on the substrate with random distribution (Supplementary Fig. 4). When the NM was placed on the edge region of the top surface of the circular magnet (Fig. 1f), the magnetic field showed a distinct gradient distribution (Fig. 1g). In this condition, the size of the prototissues in the NM decreased gradually from the center to the edge regions of the circular magnet (Fig. 1h). The red dashed section line conformed the gradient distribution of the prototissues in Figure. 1h. More importantly, this method was used to fabricate spatial programable multi component prototissues by varying the spatial magnetic field and the addition order of different GUVs (or cells).

### Diverse multi-component prototissues

The biological tissue functions depends on diverse tissue structures and heterogeneous compositions^4^. Herein, rGUVs (red fluorescence), gGUVs (green fluorescence) and non-labeled GUVs were used as building blocks to generate multi-component prototissues with diverse structures. Spatial programmable prototissues can be obtained by tuning the addition order/number of different GUVs and the distribution of magnetic fields. With a vertical field, the prototissues with rGUVs inside gGUVs colonies (Fig. 2a) was fabricated by adding gGUVs (3×10^5^/mL) and rGUVs (2×10^5^/mL) successively. With a vertical field for gGUVs (3×10^5^/mL) assembly, while with an inclined field for rGUVs (1×10^5^/mL), the rGUVs was controlled to locate at the edge of half ‘green eyes’ (Fig. 2b). With both inclined field for successive addition of gGUVs (1×10^5^/mL) and rGUVs (1×10^5^/mL), the prototissues inside each grid were shown in Figure 2c and Supplementary Figure 5. Thus, we demonstrated that the spatial positions of components were controlled by adjusting the magnetic field and the addition order/number of components.

**Fig. 2.**
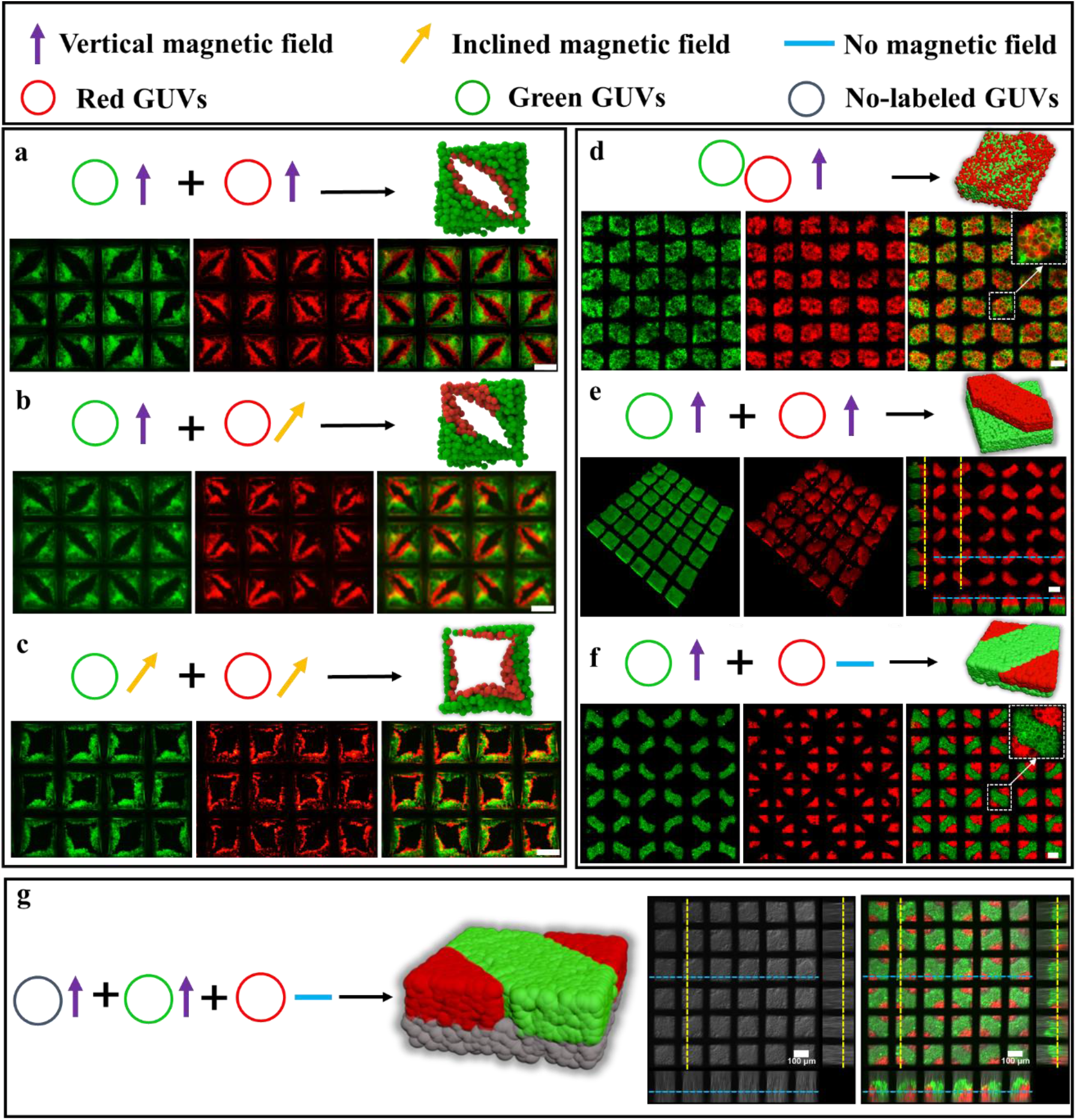
Diverse multi-component prototissues. **a)** Schematic and fluorescence images of a GUVs prototissue of ‘eye’ structures with rGUVs inside gGUVs under a vertical magnetic field with the addition of gGUVs (3×10^5^/mL) and rGUVs successively (2×10^5^/mL). (**b)** Schematic and fluorescence images of a GUVs prototissues of modified ‘eye’ structures by trapping successively gGUVs (3×10^5^/mL) under a vertical magnetic field and rGUVs (1×10^5^/mL) under an inclined magnetic field. **c)** Schematic and fluorescence images of prototissues by trapping successively gGUVs (1×10^5^/mL) and rGUVs (1×10^5^/mL) under an inclined magnetic field. **d)** Schematic and fluorescence images of binary prototissues of ‘protruded structures’ by trapping the mixture of gGUVs (6×10^5^/mL) and rGUVs (6×10^5^/mL) under a vertical magnetic field. **e)** Schematic and fluorescence images of prototissues of ‘protruded structures’ with gGUVs at the bottom and rGUVs at the top by trapping successively gGUVs (6×10^5^/mL) and rGUVs (4×10^5^/mL) under vertical magnetic field. **f)** Schematic and fluorescence images of prototissues with two layered structures by trapping successively gGUVs (1.2×10^6^/mL) under a vertical magnetic field and rGUVs (4×10^5^/mL) in the absence of magnetic field. **g)** Schematic and fluorescence images of prototissues composed of three components by trapping successively non-labeled GUVs (6×10^5^/mL) and gGUVs (6×10^5^/mL) under vertical magnetic field to form ‘protruded structrues’, and subsequently rGUVs (2×10^5^/mL) in the absence of magnetic field. All the prototissues were assembled on the top of a circular magnet with 0.3 T magnetic flux density. After one type of GUVs were trapped, the time intervals were 1 hour before adding another type of GUVs. The scale bars were 100 μm.

With the addition of more vesicles, the two layered two-component prototissues were fabricated. With a vertical field, the structures similar to Figure 1d IV but composed of rGUVs (6×10^5^/mL) and gGUVs (6×10^5^/mL) were obtained with the addition of the mixture of rGUVs and gGUVs (Fig. 2d). The green and red GUVs were evenly mixed in the prototissues. With a vertical field, the ‘protruded structures’ (Fig. 2e) with green bottom layer and red top layer were obtained by adding gGUVs (6×10^5^/mL) and rGUVs (4×10^5^/mL) successively. The projected images in Figure 2e (right image) clearly showed the protruded two layered structures. With a vertical field, the ‘protruded structures’ were formed with gGUVs (1.2×10^6^/mL) first. Subsequently with the addition of rGUVs (4×10^5^/mL) with no magnetic field, the rGUVs filled in the rest regions of top layer to form the structures as shown in Figure 2f.

More diversely, the three components spatial coded prototissues (Fig. 2g) were obtained with the addition of non-labelled GUVs (6×10^5^/mL) in a vertical field first, green GUVs in a vertical field second (6×10^5^/mL), and red GUVs with no magnetic field finally (2×10^5^/mL). The projected images in Figure 2g (right image) clearly showed the wanted two layered structures. The ratio of different GUVs can be adjusted freely to form similar structures (Supplementary Fig. 6). Based on the principle demonstrated above, more complicated programmed spatial coded prototissues can be obtained by varying the magnetic field and the addition order/number of GUVs, which makes great sense in building complicated artificial or living tissues.

### Signal communication between two-component spatial coded prototissues

In living systems, tissues are composed of heterogeneous cell populations with spatial distributions. Their functions are controlled by the signal communications among cells. After demonstrating the spatial coded prototissues with multi-components, the signal communications among them were investigated. The prototissues similar to those in Figure 2f were prepared using green GOx-gGUVs (with melittin pores in the lipid bilayer membrane, and glucose oxidases inside GUVs) and non-labeled HRP-GUVs (with horseradish peroxidases inside) (Fig. 3). The GOx-gGUVs formed the protruded structure at a vertical magnetic field, while the HRP-GUVs occupied the rest top layer regions with no magnetic field, as shown in the left image of Figure 3b. The green fluorescent image (left image in Figure 3c) and the merged image of green and bright field image (right image in Figure 3c) of the produced prototissues confirmed the designed structures. The signal communication between these two type GUVs were demonstrated using the cascade enzyme reactions shown in Figure 3a. The added glucose molecules (30 mM) entered into GOx-gGUVs via melittin pores in the bilayer membrane, subsequently to be oxidized by GOx to produce H_2_O_2_. H_2_O_2_ diffused into nearby non-labeled HRP-GUVs to oxidase Amplex Red (0.05 μM) to generate red fluorescent resorufin catalyzed by HRP. As the reactants entered into the GUVs continuously, the red fluorescence gradually became stronger and reached the equilibrium in the non-labeled GUVs regions at about 20 minutes (Fig. 3d, e). Using GOx-free gGUVs to replace the GOx-gGUVs in the structures, no red fluorescent resorufin was observed (Fig. 3e, black line). Thus, we demonstrated the signal communication between two-component spatial coded prototissues.

**Fig. 3.**
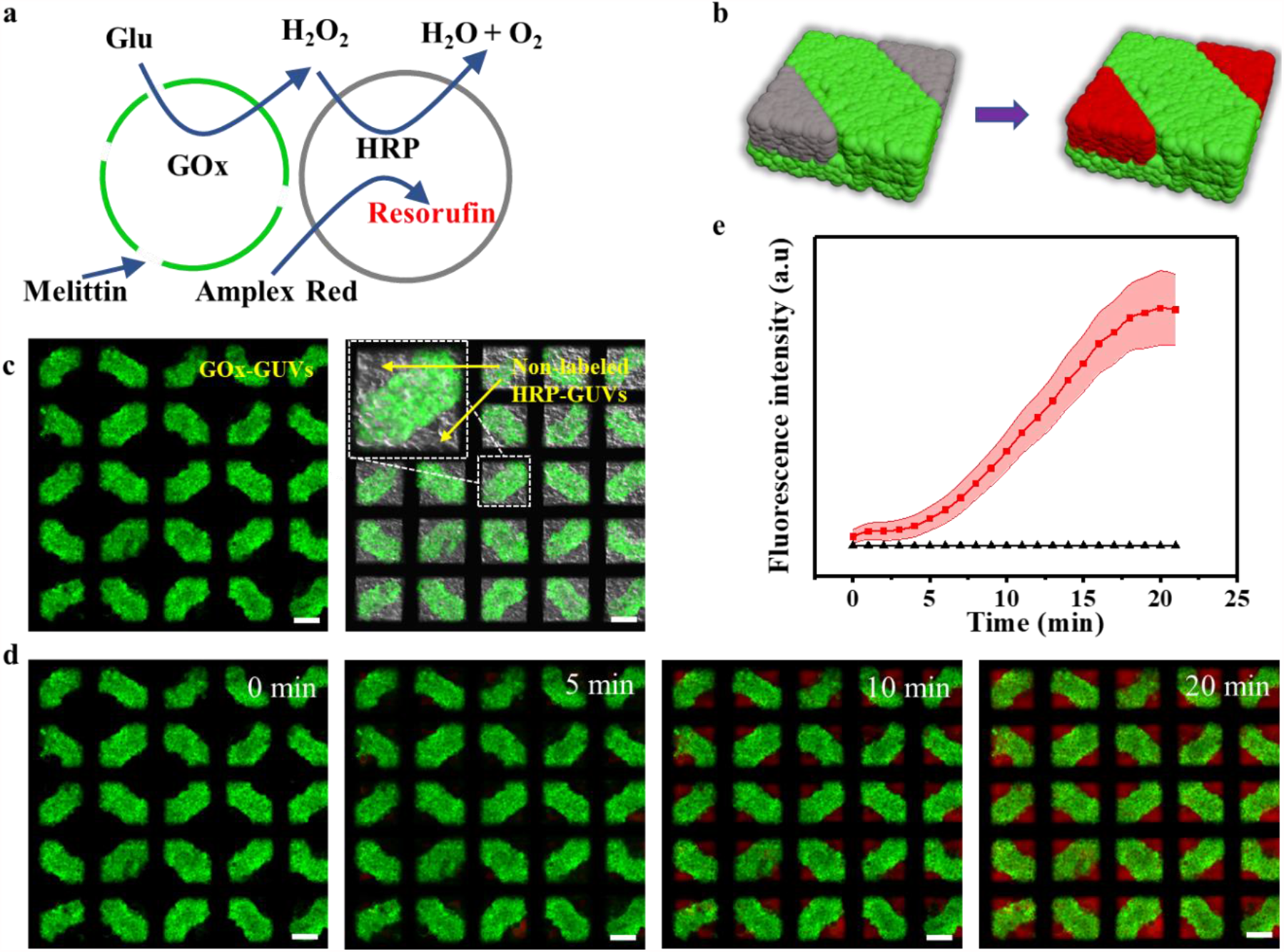
Signal communication between two-component spatial coded prototissues. **a, b)** Schematic illustration of signal communication in the prototissues composed of GOx-gGUVs with melittin pores and non-labeled HRP-GUVs. **c)** Fluorescence image of the GOx-gGUVs populations (left), the merged image of the fluorescence and bright field images of the GOx-gGUVs and non-labeled HRP-GUVs populations. **d)** Confocal fluorescence images of the prototissues as a function of time after the addition of 30 mM glucose and 0.05 μM Amplex Red. **e)** The mean fluorescence intensities in the HRP-GUVs regions in the images in d (red line), and the control samples using gGUVs to replace GOx-gGUVs (black line). The error bars were the standard error of mean (SEM, n=5). The scale bars were 100 μm.

### Signal communication in three-component spatial coded prototissues

Furthermore, the signal communications among three-component prototissues were demonstrated as schematically shown in Figure 4a and b. The protocol for fabricating the prototissues in Figure 2g was used to form the prototissues composed of three components. The bottom layer and protruded layer were composed of non-labeled GOx-GUVs and C6 glioma cells, respectively. The rest regions of top layer were occupied with Arginine-rGUVs (red TR DHPE labeled GUVs containing 20 mM L-Arginine). To initiate the ternary signal communications, the added glucose (30 mM) entered the non-labeled GOx-GUVs from the melittin pores to generate H_2_O_2_ by GOx (Fig. 4b, left image), which entered into the Arginine-rGUVs and reacted with L-Arginine to release nitric oxide (NO)^24,25^. NO then entered the living C6 glioma cell populations and interacted with NO fluorescent probe of DAF-FM dyes in cells, which generated highly fluorescent triazole derivatives (DAF-T) (Fig. 4b, right image)^26^. The experimental results (Fig. 4c) confirmed the abovementioned signal communication pathway. The protruded C6 cell regions exhibited weak green fluorescence at 0 hour, but gradually stronger green fluorescence as a function of time (Fig. 4c). The corresponding fluorescence intensity curve was shown in Figure 4d (green curve). There was almost no increase of the fluorescence signal when GOx-GUVs were replaced by GOx-free GUVs (Fig. 4d, red curve). However, when L-Arginine was absent in rGUVs, the fluorescence intensity showed a low-level increase, which was probably due to the inherent L-Arginine in cells interacting with H_2_O_2_ (Fig. 4d, blue curve). A confocal fluorescence image with projected images of prototissues composed of three components showed the positions of non-labeled GOx-GVUs populations (the gray areas) at the bottom layer, Arginine-rGUVs populations (the red areas) at the edge of top layer, and cell populations (the green areas) at the top layer (Fig. 4e and Supplementary Fig. 7).

**Fig 4.**
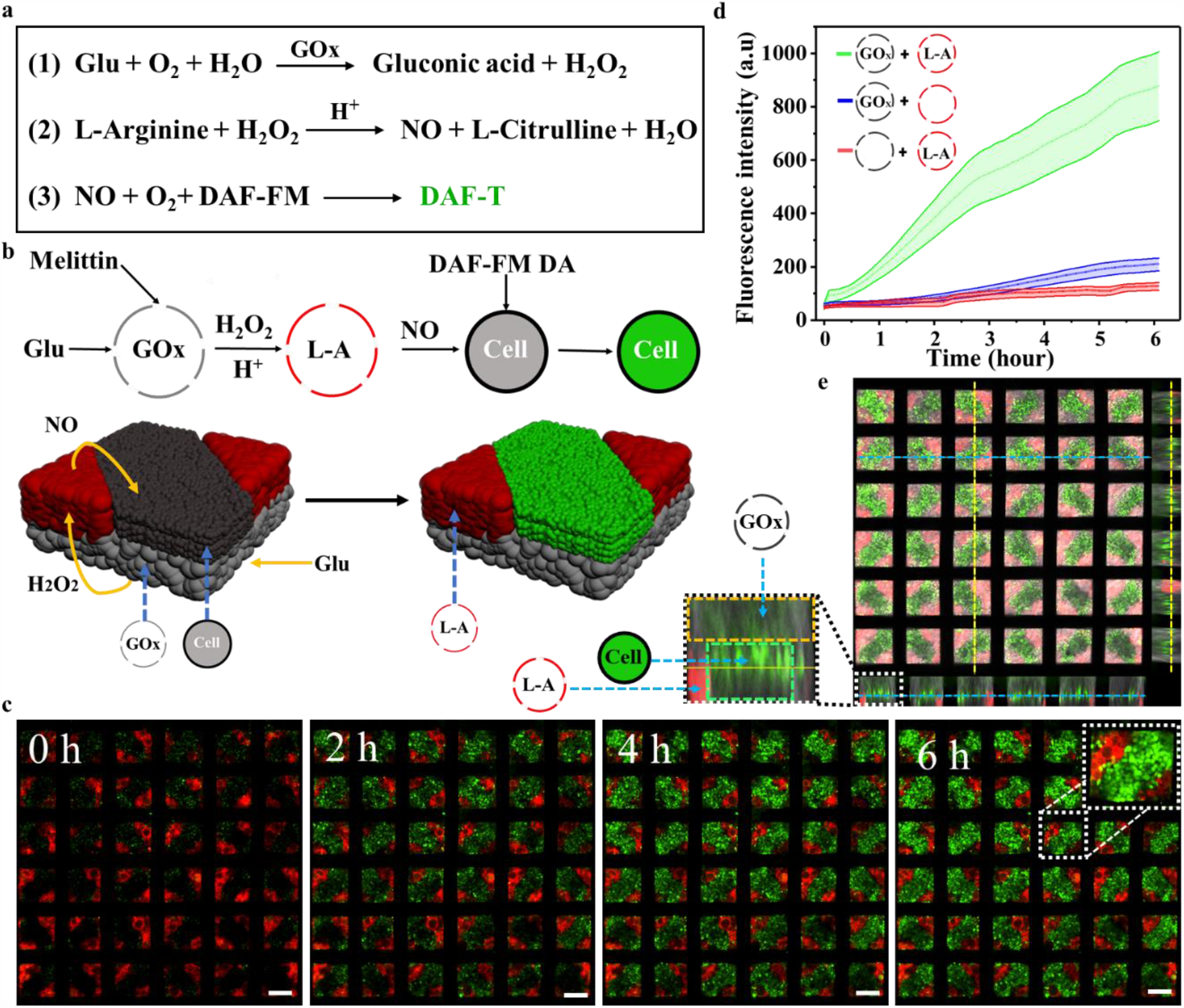
Signal communications in hybrid three-component prototissues composed GUVs and C6 glioma cells. **a)** The cascade reaction formulas for the ternary signal communications among prototissues. **b)** Schematic illustration of signal communications among these three components in the prototissues with GOx-GUVs at the bottom, C6 glioma cells at the protruded top layer, and Arginine-rGUVs (containing 20 mM L-Arginine (L-A)) at the rest regions of top layer. **c)** Fluorescence images of the prototissues as a function of time triggered by the addition of glucose (30 mM) in the solution. The red regions indicated the Arginine-rGUVs populations. The green regions indicated the cell populations. **d)** The corresponding green fluorescence intensities of prototissues in **c** (green line), same prototissues but using Arginine-free GUVs to replace Arginine-rGUVs (blue line), and the same prototissues but using GUVs with no GOx to replace GOx-GVU_s_ (red line). The error bars were the standard error of mean (n=5). **e)** A confocal fluorescence image with projected images of the hybrid prototissues composed of three components at 6 hours after the addition of glucose. The gray, red and green regions indicated the non-labeled GOx-GUVs bottom layer, the Arginine-rGUVs top edge layer, and the protruded cell top layer. The scale bars were 100 μm.

### Prototissues capable of producing NO

The functions of prepared prototissues triggered by the internal signal communications were investigated in the following contents. Given NO served commonly as a second messenger involved in many physiological functions, such as inducing the relaxation of blood vessels^27,28^. The prototissues were detached from the NM grids by shaking for the potential biomedical application. They still kept their morphology because of the hemi-fusion among gGUVs by incubating in 100 mM CaCl_2_ for ten minutes before they were detached from NM grids (Fig. 5a). They allowed the molecule smaller than 20 kDa to enter the interior of prototissues to some extent (Fig. 5b). The signal communications among the detached prototissues composed of two components (Fig. 5c, d) were investigated to produce NO. It is well known that hydroxyurea (Ha) is oxidized by H_2_O_2_ to generate NO in the presence of HRP ^29^. The prototissues composed of non-labeled GOx-GUVs with melittin pores and HRP-rGUVs were fabricated (Fig. 5d, bottom). After the addition of hydroxyurea (10 mM) and glucose (20 mM), a rapid increasing green fluorescence in the prototissues was observed due to the fluorescence product of DAF-2T generated by the interaction of NO and DAF-2 (10 μM) in the solution (Fig. 5e, pink line in f). As the diffusion of NO from prototissue, the fluorescence intensity in exterior region was increasing against time (Fig. 5e, blue line in f), which confirmed the NO-generating capacity of the prepared prototissues.

**Fig 5.**
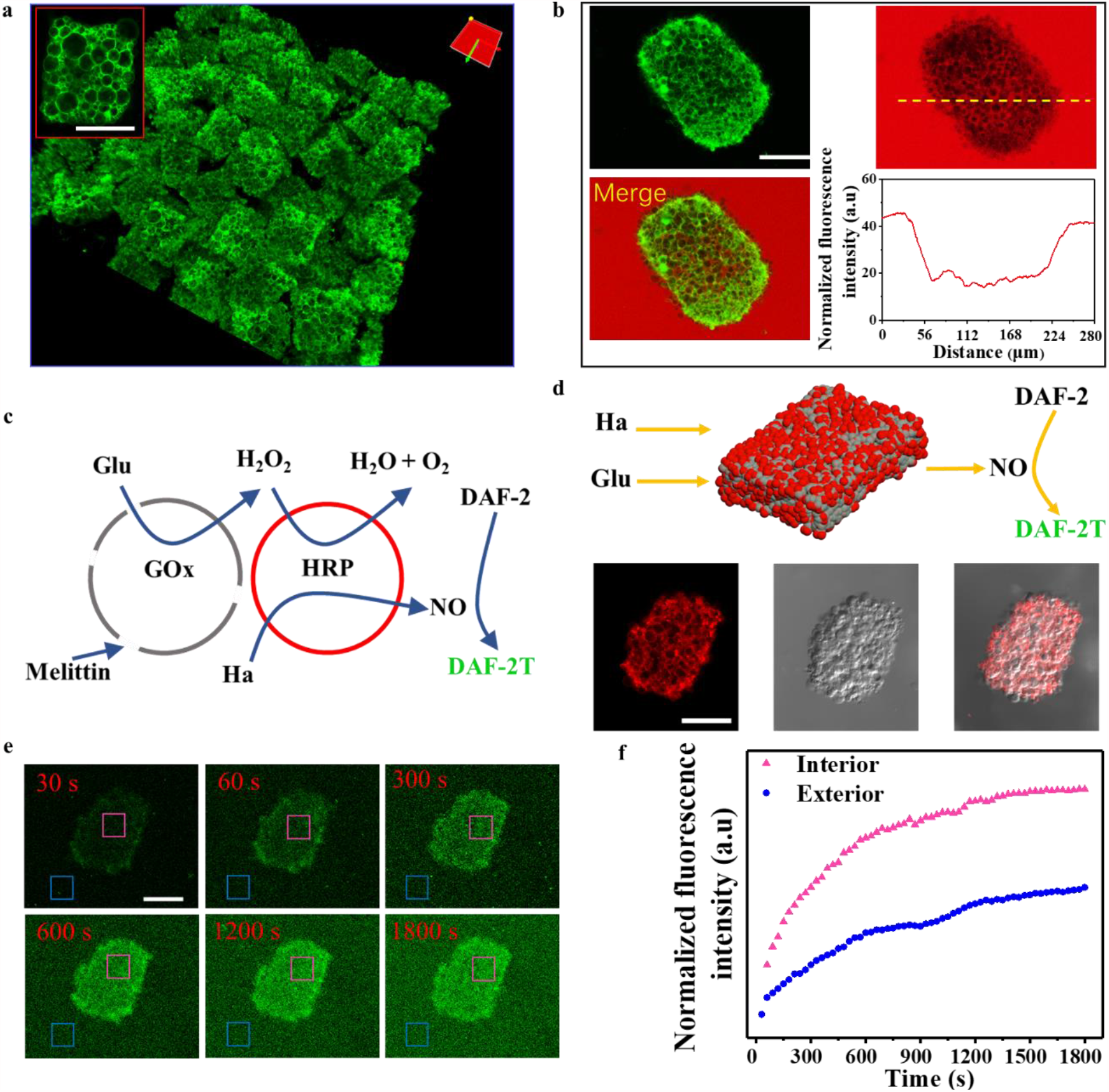
Prototissues capable of producing NO. **a)** The 3D reconstructed confocal image of the GUVs prototissues detached from the NM. The red box indicated an enlarged GUVs prototissue. **b)** Fluorescence images of the detached prototissue in Rhodamine-dextran (20 kDa) solution. The yellow dashed line in the fluorescence image correspond to line intensity analysis. **c)** Schematic illustration of signal communcations among NO-prototissue composed of GOx-GUVs with melittin pores and HRP-rGUVs. The addition of glucose and hydroxyurea triggered the reaction to generate NO in HRP-rGUVs. NO was detected by the DAF-2 in the solution to generate green fluorescent DAF-2T. **d)** Schematic illustration of the structure of the NO-prototissue (top). Fluorescence image of the HRP-rGUVs in the NO-prototissue (bottom left), bright field image (bottom middle) and their merged image (bottom righ) of the NO-prototissue. **e)** Time-dependent fluorescence microscopy images of the NO-prototissue in the solution containing DAF-2 (10 μM) after adding glucose (20 mM) and hydroxyurea (10 mM). The green fluorescence channel responded to NO production. **f)** Plots of fluorescence intensity against time for internal (pink box) and external regions (blue box) in **e)**. The scale bars were 100 μm.

### NO-prototissues for vasodilation

The prototissues capable of producing NO (NO-prototissues) were further investigated with living blood vessel tissue to prove their potential biomedical applications. The NO-prototissues were mixed with living blood vessels. With the addition of hydroxyurea, NO-prototissues produce NO molecules, which cause blood vessel relaxation (Fig. 6a). A thoracic aorta ring test was used to validate the function of NO-prototissues. A 3 mm wide thoracic aorta ring was hung on the hooks connected to a force transducer in the HEPEs solution containing 20 mM glucose. Upon the addition of hydroxyurea (10 mM) at 10 minute, NO-mediated vasodilation was immediately reflected by the force sensor, which exhibited a rapid decrease of tension force (Fig. 6b, left). For a comparison, there was no obvious change of the tension force when NO-free-prototissues composed of GOx-free GUVs and HRP-GUVs were added into the solution at 15 minutes (Fig. 6b, right), because NO-free-prototissues did not produce NO molecures. The variation of the tension force were calculated before and 5 minutes after the addition of hydroxyurea. Typically, NO-prototissues resulted in a tension force decrease of 2.59 g and a blood vessel relaxation of 31.2% (Fig. 6c, d). When the blood vessels were treated by the NO-free-prototissues, there was a small tension force decrease of 0.14 g and a blood vessel relaxation of 3.4%, which was attributed to the low-level degradation of hydroxyurea in the media (Fig. 6c, d). Moreover, the sequential addition of the same amount of hydroxyurea at 30 minutes (Fig. 6b, left) induced the blood vessels to relax again, which confirmed that the blood vessels remained viable within the experimental period. Fluorescence images of blood vessels sections also confirmed the formation of NO in the vascular ring through a cell-permeable NO dye (DAF-FM DA) (Fig. 6e). NO-prototissues induced high intensity NO-mediated green fluorescence in blood vessel (Fig. 6e, top row). On the contrary, lower fluorescence intensity was observed in NO-free-prototissues mediated experiments due to the small amount of degraded hydroxyurea (Fig. 6e, bottom row). These experimental results implied the potential application of NO-prototissue in cardiovascular disease by transplantations. In summary, the phospholipid vesicles and c6 glioma cells were used as the building blocks to precisely fabricate high-throughput spatial coded multi-component prototissues using the magneto-Archimedes effect. The signal communications were demonstrated in the spatially programmable two- and three-component prototissues using two-step and three-step enzyme cascade reactions, respectively. Significantly, the prototissues composed of GOx-GUVs and HRP-GUVs produced NO via the internal communications in the prototissues triggered by glucose and hydroxyurea. These prototissues were confirmed to be capable of relaxing the mouse blood vessels, consequently to improve their functions, which may be used to treat cardiovascular diseases through transplantations.

**Fig 6.**
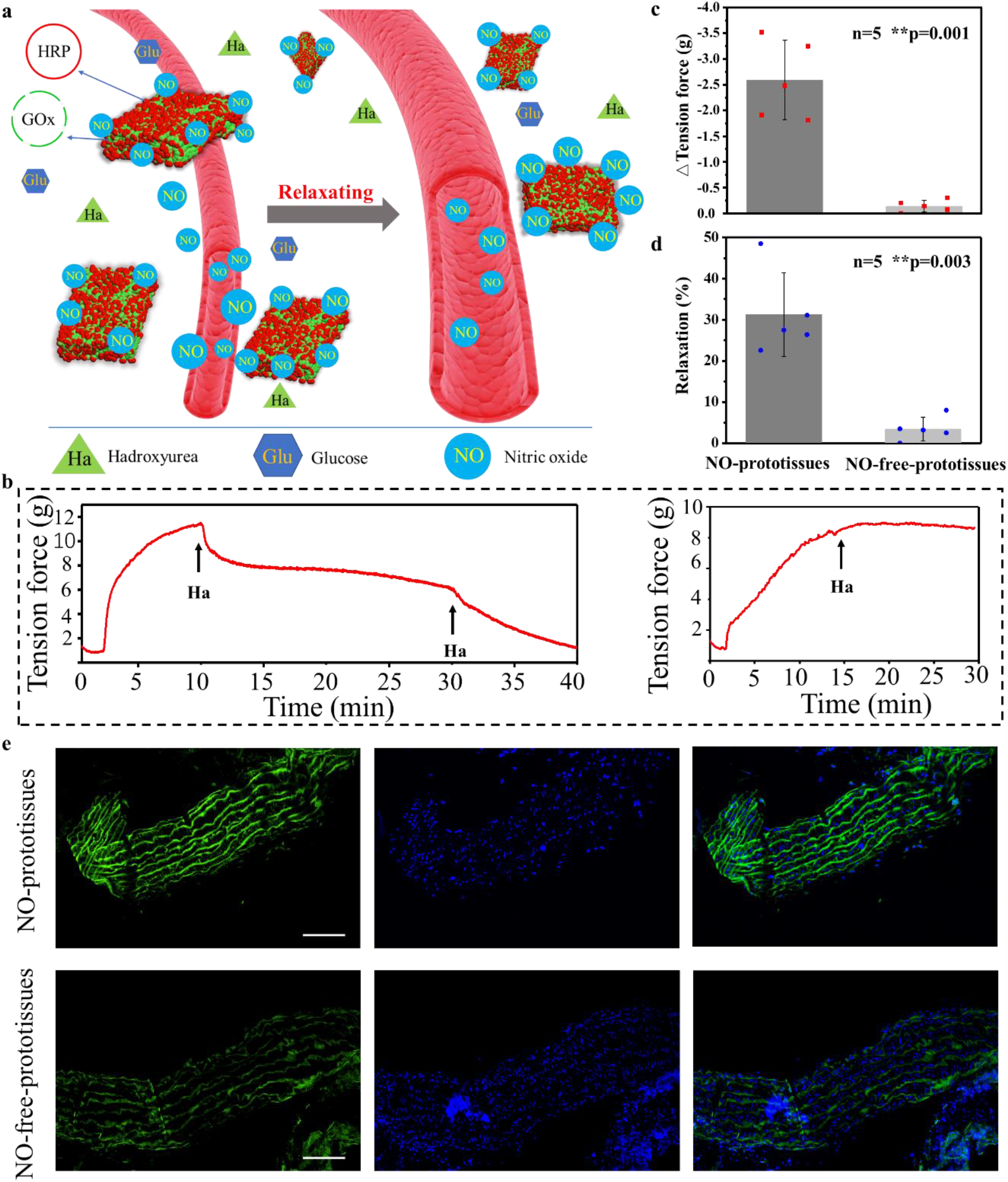
NO-prototissues for vasodilation. **a)** Schematic illustration of vasodilation induced by NO from NO-prototissues. **b)** Representative tension curve of vasodilation against time in the presence of NO-prototissues composed with GOx-GUVs and HRP-GUVs (left), and NO-free-prototissues composed of GOx-free GUVs and HRP-GUVs (right). The black arrow indicated the point of the addition of hydroxyurea (Ha, 10 mM). **c), d)** Bar charts of the decrease in tension force and relaxation of vascular rings observed in the presence of NO-prototissues (2.59 g; 31.2% relaxation) and NO-free-prototissues (0.14 g; 3.4% relaxation), n=5. **e)** Fluorescence images of vascular sections stained with DAF-FM DA (the green fluorescence channel responded to NO production, left column), DAPI (blue fluorescence with nuclei, middle column), and their merge images (right column). The blood vessels were treated using NO-prototissues (top row) or NO-free-prototissues (bottom row) in the HEPES solution containing 10 mM hydroxyurea for 20 minutes, respectively. The scale bars were 100 μm.

## Methods

### Materials

1,2-Dioleoyl-sn-glycero-3-phosphocholine (DOPC), Texas Red-labeled 1,2-dihexadecanoyl-sn-glycero-3-phosphoethanolamine triethylammonium salt (TR-DHPE) and N-(7-nitrobenz-2-oxa-1,3-diazol-4-yl)-1,2-dihexadecanoyl-sn-glycero-3-phosphoethanolamine triethylammonium salt (NBD-PE) were purchased from Avanti Polar Lipids (USA). Horseradish peroxidase (HRP), glucose oxidase (GOx), amplex red, melittin, hydroxyurea (Ha), 4-aminomethyl-2’,7’ - difluorofluorescein diacetate (DAF-FM DA), 2-(3,6-dihydroxy-4,5-diamino-9H-xanthen-9-yl)-benzoic acid (DAF-2), DAPI and L-Arginine (L-A) were purchased from Sigma (USA). Sucrose (Suc), glucose (Glu), Manganese (II) (MnCl_2_) chloride, Calcium chloride (CaCl_2_) and hydrochloric acid (HCl) were obtained from Aladdin (China). Gadobutrol injections were obtained from Harbin Institute of Technology Hospital (China). The nickel meshes (NM) were purchased from Gates RGRS Company (China). Cylindrical NdFeb magnets (1 T, diameter = 3 cm, thickness = 1 cm) were bought from Gates Qiangci Company (China). The Indium tin oxide (ITO) electrodes were obtained from Hangzhou Yuhong technology Co. Ltd (China). Dulbecco’s modified Eagle’s medium (DMEM), phosphate buffer saline (PBS) without calcium and magnesium, trypsin, streptomycin and penicillin were purchased from Corning (USA). The fatal bovine serum (FBS) were purchased from Gibco (USA).

### Preparation of GUVs

The formation of giant unilamellar vesicles (GUVs) was described elsewhere (Supplementary Fig. 8)^30-32^. In brief, DOPC (5 mg/mL, 8 μL) mixed with the fluorescent lipids TR-DHPE (0.5% w/w, red fluorescence) or NBD-PE (5% w/w, green fluorescence) in chloroform was gently spread on two ITO electrodes to form lipid films. A rectangular polytetrafluoroethylene frame was placed between two lipid film coated ITO electrodes to form the setup. An AC electric field (5 V, 10 Hz) was applied by a signal generator for 1 h, followed by applying an electric field (0.8 V, 2 Hz) for additional 5 min. Different solutions were filled inside the frame to prepare the wanted GUVs. The sucrose solution (300 mM) containing 20 μg/mL HRP or 30 μg/mL GOx was used to obtain HRP-GUVs or GOx-GUVs. The sucrose solution (300 mM) containing 20 mM L-Arginine was used to prepare Arginine-GUVs. The free GOx, HRP and L-Arginine in the supernatants were removed by centrifugation (2000 r, 5 min, 5 times), until no corresponding molecules were detected in the supernatants (Supplementary Fig. 9). The GUVs were incubated in solution containing 50 μg/mL melittin for 20 mins to obtain the GUVs with melittin pores in the lipid bilayer membrane.

### Formation of the prototissues

Prototissues were prepared using a home-made device (Supplementary Fig. 2). A square NM (1.2 cm × 1.2 cm) was fixed in a petri dish using vacuum grease. To avoid GUVs rupture during assembly, 200 μL of 0.1 mg/ml DOPC ethanol-water solution with ethanol volume percentage of 40% was added in the petri dish at 45 °C for 10 minutes, followed by washing the device with MnCl_2_ solution or PBS for three times. For the formation of prototissues under vertical magnetic field, the device was placed on the center of the top surface of a circular permanent magnet, followed by adding GUVs labeled with NBD-PE (gGUVs) or TR DHPE (rGUVs) in 100 mM MnCl_2_ solution. The inclined magnetic field was provided by putting the NM at the edge regions of the top surface of the magnet as shown in Fig. 1f. For the formation of two-component prototissues (Fig. 2a, b, c, e and f), gGUVs and rGUVs were successively added into the device under the vertical or inclined magnetic field. The two-component prototissues in Fig. 2d was obtained by adding the mixture solution of gGUVs and rGUVs into the device under the vertical magnetic field. For the formation of three-component prototissues in Fig. 2g, non-labeled GUVs and gGUVs were successively trapped under the vertical magnetic field, and then rGUVs were trapped in the absence of the magnetic field. After one type of GUVs were trapped, the time intervals were 1 hour before adding another type of GUVs. The concentration of GUVs was controlled by varying the volume ratio of the GUVs sucrose solution and MnCl_2_ (gadobutrol solution). Before preparing prototissue array containing living cells, the device and the solution were sterilized before use and the MnCl_2_ solution was replaced by PBS solution containing 30 mM gadobutrol. The device was placed at 37 °C and 5% CO_2_ in a humidified atmosphere throughout the experiments containing cells.

### Cell line and cell culture

C6 glioma cells were cultured in DMEM containing 50 μg/mL penicillin, 50 μg/mL streptomycin and 10% fetal bovine serum at 37 °C and 5% CO_2_ atmosphere. The cells were digested by trypsin, concentrated by centrifugation at 2000 rpm for 5 min, and re-dispersed in the medium with the final density of 2×10^5^/Ml.

### Signal communications in prototissues

The GOx-gGUVs with melittin pores and HRP-GUVs were magnetically assembled into the structure as shown in Figure 3b. To initiate the signal communication, the external solution was replaced with 30 mM glucose containing 0.05 μM Amplex Red. The fluorescent product was monitored using fluorescence microscope. For comparison, GOx-free gGUVs were used to replace the GOx-gGUVs in the prototissues. To form the prototissue as shown in Figure 4b, the GOx-GUVs with melittin pores and C6 glioma cells were successively trapped under the vertical magnetic field, and then Arginine-rGUVs with melittin pores (containing 20 mM L-Arginine) were trapped without magnetic field in the PBS containing 30 mM gadobutrol. C6 glioma cells were incubated in PBS containing NO probe (10 μM DAF-FM DA) for 20 minutes before adding them into the device. To initiate the signal communication, the external solution was replaced with PBS containing 30 mM glucose. For comparisons, GOx-GUVs or Arginine-rGUVs in the prototissue were replaced by GOx-free GUVs or Arginine-free rGUVs, respectively.

### Detachment of prototissues capable of producing NO

After the GUVs prototissue array was assembled on the NM, 100 mM CaCl_2_ was used to promote hemi-fusion of the prototissues composed of no-labeled GOx-GUVs with melittin pores and HRP-rGUVs for 10 minutes. The prototissues were detached from the NM by gently shaking in the solution. We named these prototissues as NO-prototissues. Afterwards, the prototissues were washed 3 times using PBS to remove CaCl_2_ in the solution. The permeability of prototissue was detected using Rhodamine-dextran (20 kDa). 10 mM hydroxyurea and 20 mM glucose were added into the device to trigger the reaction. 10 μM DAF-2 was used to detected the released NO from prototissues.

### Vasodilation stimulated by NO from NO-prototissues

The thoracic aortas of mice were separated after the mice were anesthetized (n = 5 vascular rings per group, 10 vascular rings, 5 mice). The blood vessels were cut into rings after the extraneous fat and connective tissues were removed mechanically. The rings were connected to a force transducer in the baths containing 10 mL of HEPEs solution (component (mM): NaCl 137.6, KCl 4.0, MgCl_2_ 1.1, CaCl_2_ 2.1, NaHCO_3_ 10.0, NaH_2_PO4 0.92 and Glucose 20, pH 7.4). After equilibration for 20 min, a high potassium concentration solution (KCl, 70 mM) was used to induce the contraction of the vascular rings. When the tension force reached equilibrium, about 1×10^4^ NO-prototissues composed of GOx-GUVs with melittin pores and HRP-GUVs and 10 mM hydroxyurea were added into the bath solution. The tension force was detected by the force transducer. For comparison, the NO-free-prototissues composed of GOx-free GUVs and HRP-GUVs were used to replace NO-prototissues. The amount of variation in tension force (ΔTension) and the percentage of relaxation (Relaxation %) were defined as follows: ΔTension force = Fa – Fb; Relaxation % = (Fa – Fb)/ Fb × 100%. Fb and Fa represented the tension force before and after adding the prototissues and hydroxyurea, respectively. Before the vascular rings were snap-frozen and cut into slices, they were incubated for 20 min in the PBS containing 10 μM DAF-FM DA after treated with the prototissues. We confirmed that ethical approval from the Experimental Animal Ethics Committee of Harbin Institute of Technology (IACUC-2021012) was obtained prior to the study.

### Instruments

The prototissues were characterized by inverted fluorescence microscope (Olympus IX73, Japan), laser confocal microscope (Olympus FV 3000, Japan) and upright fluorescence microscope (Nikon 80i, Japan). The magnetic field was simulated using COMSOL Multiphysics 5.4 software. The AC electric field was generated with a signal generator (Agilent 33220a-001). The tension force was detected by tension sensor in a constant temperature perfusion system for isolated tissues and organs (TECHMAN HV1403, China).

## Acknowledgements

This work was supported by the National Natural Science Foundation of China (Grant No. 21929401, 21773050) and the Heilongjiang Touyan Team (HITTY-20190034).

## Author contributions

X.J.H. supervised the research. X.J.H., X.X.Z. and C.L. conceived and designed the experiments. X.X.Z., C. L., F. K. L., Y.S.R., B.Y.Y and W. M. performed experiments. X.J.H., X.X.Z., C. L., F. K. L., Y.S.R., B.Y.Y and W. M. analyzed the data. X.J.H., X. X. Z., and C.L wrote the paper. All authors commented on the paper. X. X. Z and C. L contributed equally to this work.

