## supplemental information for "High-throughput production of functional prototissues capable of producing NO for vasodilation"

Han et al.

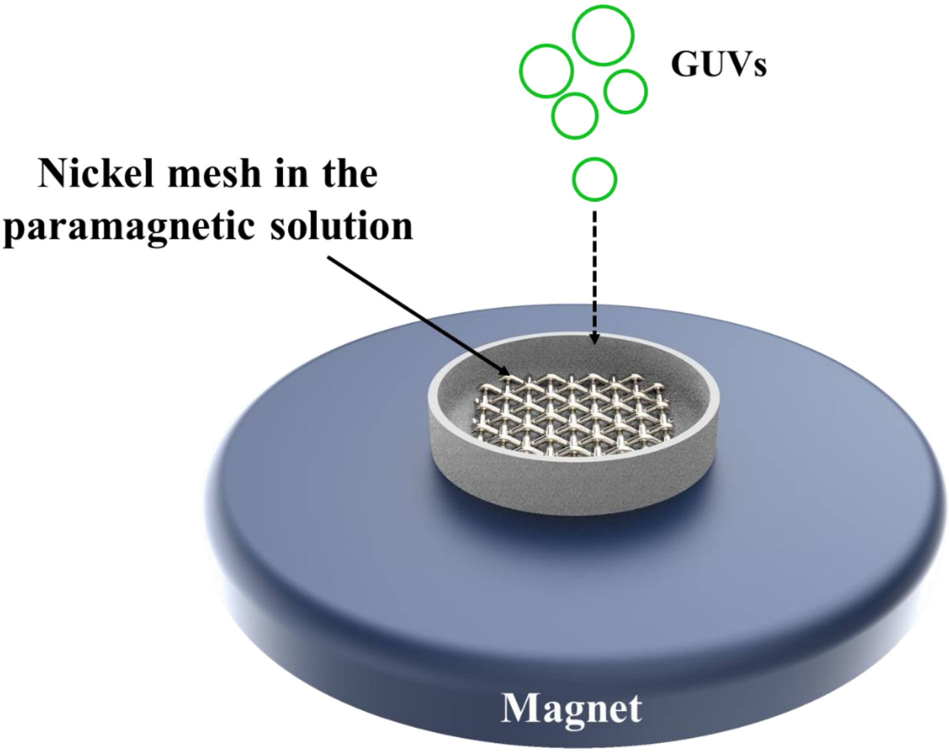


**Supplementary Fig. 1** Schematic illustration of home-made device for prototissue formation.

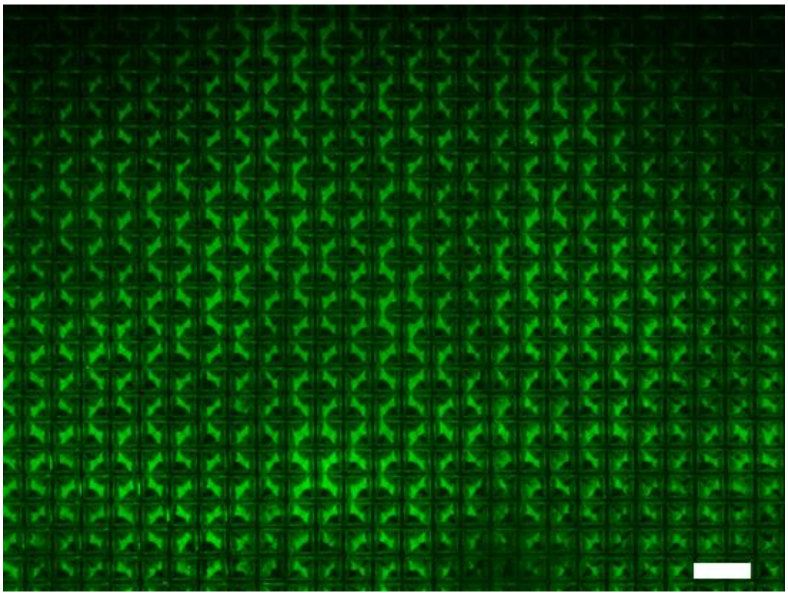

**Supplementary Fig. 2** Fluorescence zoom out image of the GUVs prototissue array. The scale bar was 400 μm.

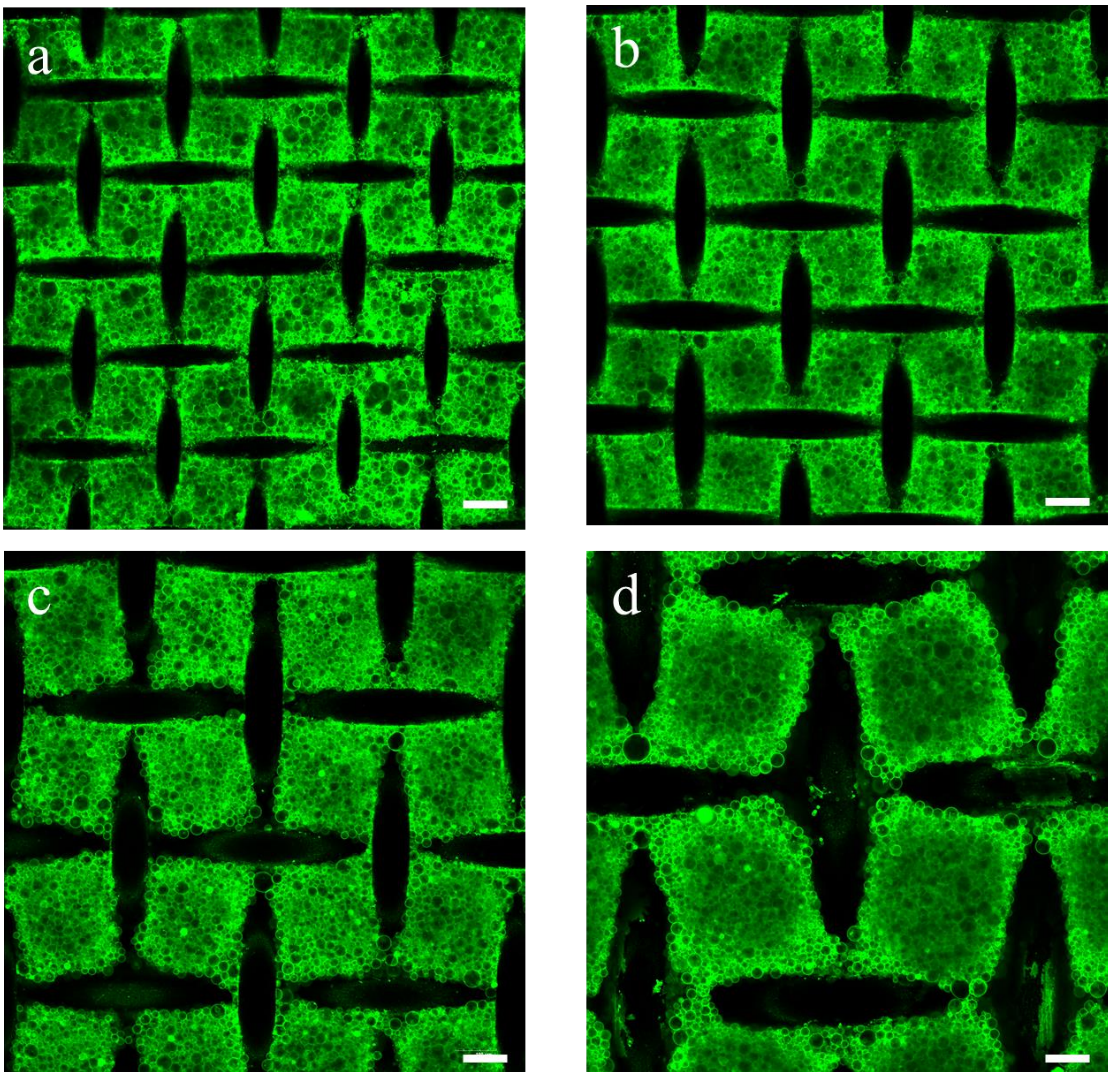


**Supplementary Fig. 3** GUVs prototissues assembled using the NMs with different side length of 150 µm (**a**), 180 µm (**b**), 230 µm (**c**), and 350 µm (d). The scale bars were 100 µm.

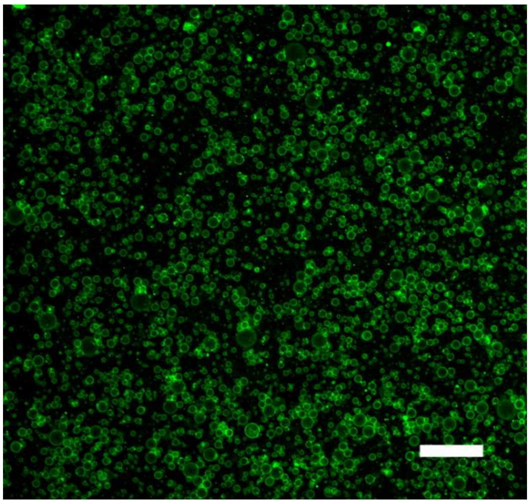

**Supplementary Fig. 4** Fluorescence image of the distribution of GUVs with no magnetic field. Random GUVs distribution was observed. The scale bar was 100 μm.

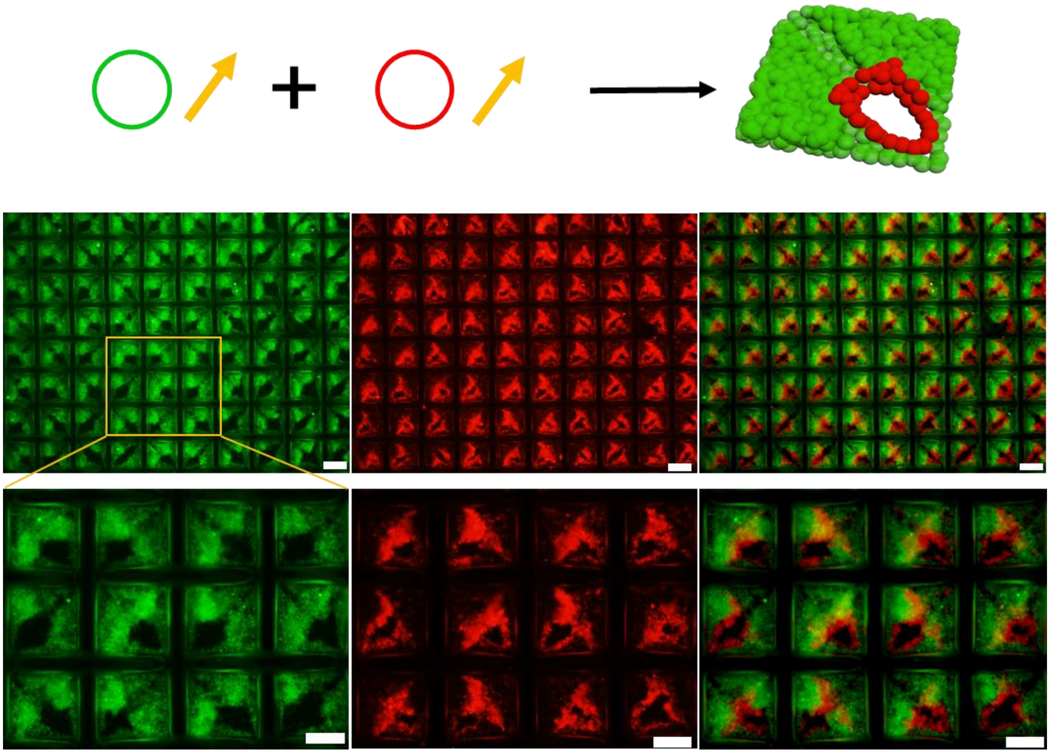

**Supplementary Fig. 5** Schematic and fluorescence images of the two-component GUVs prototissues. The prototissues inside each grid were assembled by successively trapping gGUVs ($3\times10^5$/mL) and rGUVs ($1.5\times10^5$/mL) under inclined magnetic field. The images in the bottom row were partial enlargements of the images in the top row. The left and middle column images were the prototissues viewed by green channel and red channel, respectively. The right column images were the merged images of left and middle images. The scale bars were 100 μm.

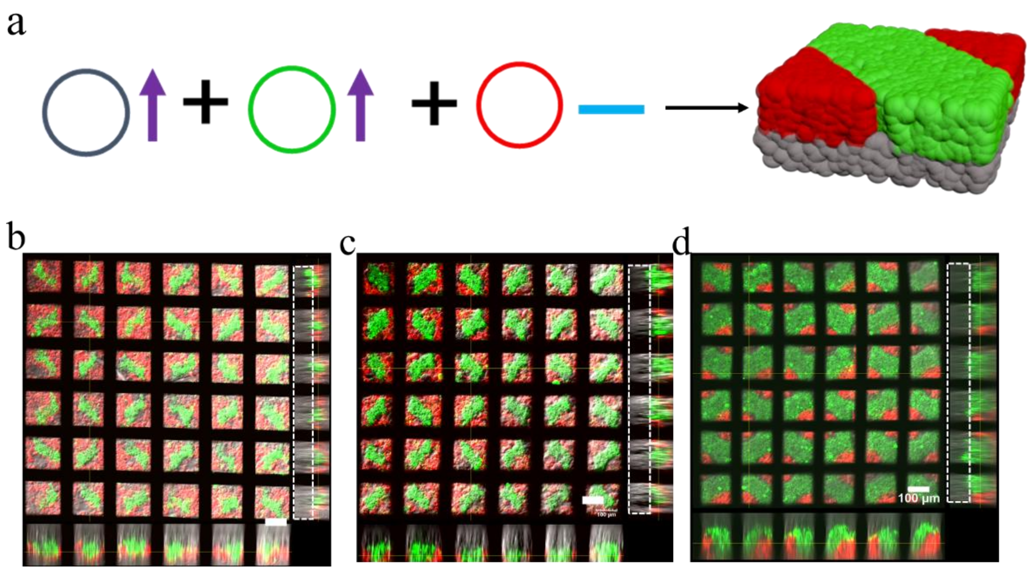


**Supplementary Fig. 6** Three-component GUVs prototissues. **a)** Schematic of the three-component prototissues assembled by successively trapping non-labeled GUVs and gGUVs under vertical magnetic field (↑), and rGUVs with no magnetic field (▬). **b, c, d)** Fluorescence images of the three-component GUVs prototissues with varying rGUVs and gGUVs. The concentrations of the non-labeled GUVs were all $6\times10^5$/mL in **b, c, d**. The concentrations of the gGUVs were $3\times10^5$/mL, $4\times10^5$/mL and $6\times10^5$/mL in **b, c, d**, respectively. The concentrations of the rGUVs were $5\times10^5$/mL, $4\times10^5$/mL and $2\times10^5$/mL in **b, c, d**, respectively. The non-labeled GUVs populations were indicated by the white dashed boxes. The scale bars were 100 μm.

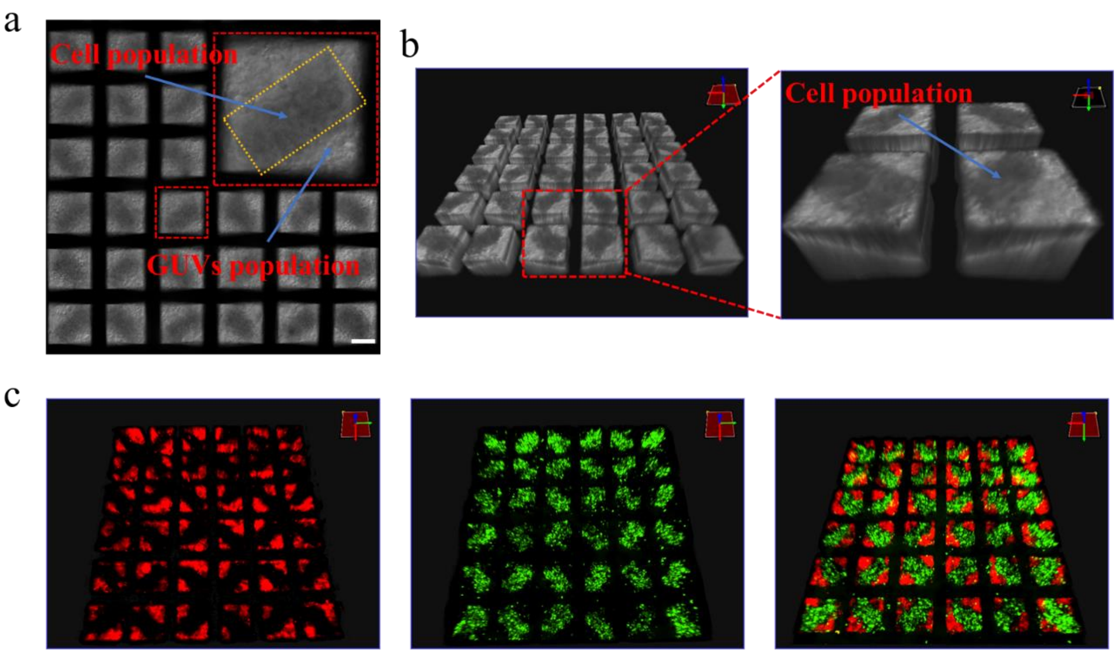


**Supplementary Fig. 7** Three-component hybrid prototissues composed of living cells and GUVs. **a)** Bright field image of the three-component prototissues. The yellow dashed box indicated the cell population. **b)** 3D reconstructed bright images of the prototissue showed the relative positions of GVUs populations and living cell populations. The black regions indicated the cell populations and the gray regions indicated the GUVs populations. **c)** 3D reconstructed confocal fluorescence images of the prototissues of rGVUs populations (left), cell populations (middle) and their merged image (right). The scale bar was 100 μm.

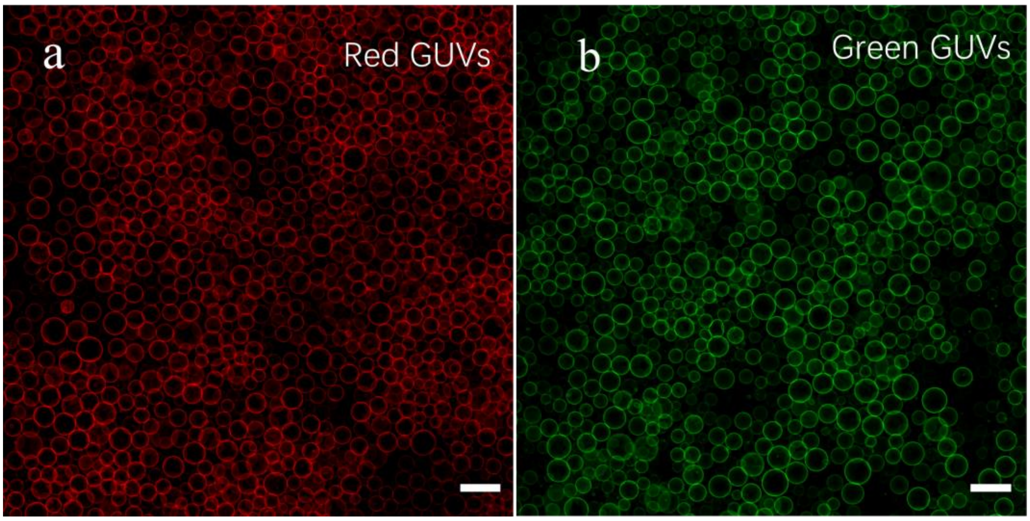


**Supplementary Fig. 8** Fluorescence images of electroformed GUVs. **a)** GUVs labelled with TR DHPE (rGUVs). **b)** GUVs labelled with NBD PE (gGUVs). The scale bars were 40 μm.

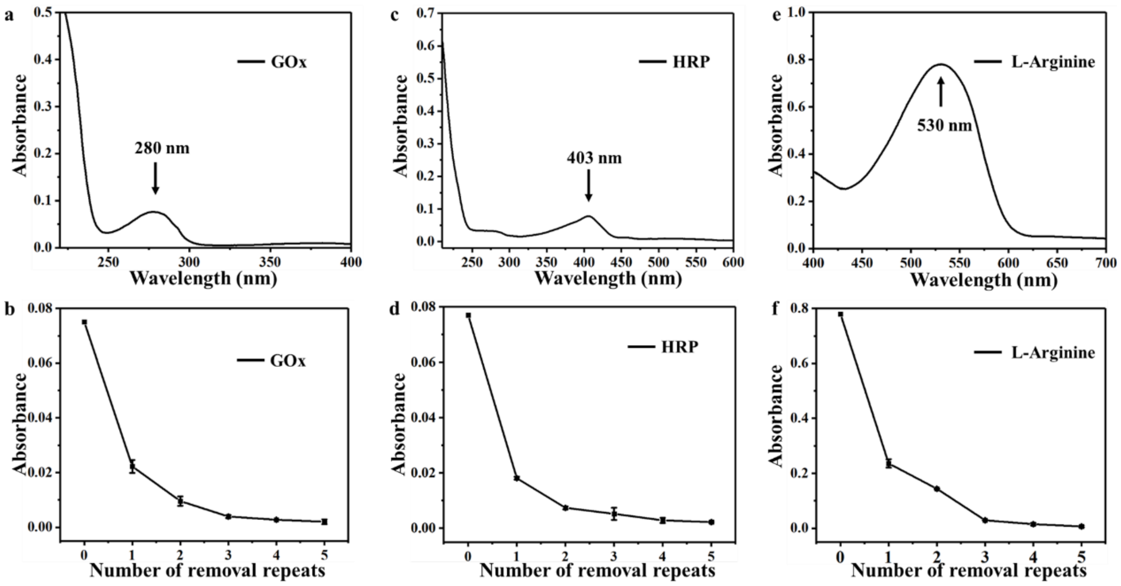


**Supplementary Fig. 9 The removal of the free GOx, HRP and L-Arginine in the supernatants. a)** Absorbance of 30 μg/ml GOx solutionor. **b)** Absorbance of supernatants of GOx-GUVs solution at 280 nm as a function of removal repeats. **c)** 20 μg/mL HRP solution. **d)** Absorbance of supernatants of HRP-GUVs solution at 403 nm as a function of removal repeats. **e)** Absorbance of 3mL 20 mM L-Arginine treated by 3.5 mL Sakaguchi's reagent[1] after 15 minutes. Sakaguchi's reagent contained 2.15 g/L NaOH solution, 42.86 g/L α-naphthol and 2.15 μL/L 2,3-butanedione. **f)** Absorbance of supernatants of Arginine-GUVs solution at 530 nm as a function of removal repeats.

### Referance

[1] Goldschmidt, M. C. & Lockhart, B. M. Simplified Rapid Procedure for Determination of Agmatine and Other Guanidino-Containing Compounds. *Analytical Chemistry*, **1971**, *43*, 1475.
